# Neural correlates of metacognition across the adult lifespan

**DOI:** 10.1101/2021.03.17.431767

**Authors:** Helen Overhoff, Yiu Hong Ko, Daniel Feuerriegel, Gereon R. Fink, Jutta Stahl, Peter H. Weiss, Stefan Bode, Eva Niessen

## Abstract

Metacognitive accuracy describes the degree of overlap between the subjective perception of one’s decision accuracy (i.e., confidence) and objectively observed performance. With older age, the need for accurate metacognitive evaluation increases; however, error detection rates typically decrease. We investigated the effect of ageing on metacognitive accuracy using event-related potentials (ERPs) reflecting error detection and confidence: the error/correct negativity (N_e/c_) and the error/correct positivity (P_e/c_). Sixty-five healthy adults (20 to 76 years) completed a complex perceptual task and provided confidence ratings. We found that metacognitive accuracy declined with age beyond the expected decline in task performance, while the adaptive adjustment of behaviour was well preserved. P_e/c_ amplitudes varied by confidence rating, but they did not mirror the reduction in metacognitive accuracy. N_e/c_ amplitudes decreased with age except for high confidence correct responses. The results suggest that age-related difficulties in metacognitive evaluation could be related to an impaired integration of decision accuracy and confidence information processing. Ultimately, training the metacognitive evaluation of fundamental decisions in older adults might constitute a promising endeavour.

## 1. Introduction

We are continuously monitoring and controlling our behaviour in order to achieve goals and avoid errors. The internal evaluation of our behaviour and our decisions, also referred to as *metacognition*, is crucial in everyday life, because it guides our present and future behaviour (Desender et al., 2018; Rabbitt, 1966). Metacognition comprises both the detection of committed errors and a feeling of confidence that accompanies a decision (Fleming and Frith, 2014; Shekhar and Rahnev, 2020). When we feel less confident about a decision, we might try to adjust it, seek more information, or recruit additional cognitive processes to optimise performance (Desender et al., 2019a, 2019b). As ageing is usually associated with declining cognitive functions and higher rates of decision errors in daily activities, decisions and corresponding motor actions need to be adjusted more often (Hertzog, 2015; Ruitenberg et al., 2014). This might be achieved, for example, by increasing efforts for an efficient metacognitive evaluation of one’s behaviour.

In general, metacognitive judgements are highly predictive of actual task performance, yet there is strong evidence that metacognition constitutes a dissociable process from the execution of the initial task (Galvin et al., 2003; Song et al., 2011). The degree to which subjective perceptions and objectively observed performance overlap, that is, the *accuracy* of metacognitive judgements, varies across individuals and task demands (Fleming & Dolan, 2012; Hertzog & Hultsch, 2000; Rahnev et al., 2020). Metacognitive accuracy has been addressed in two separate but arguably related fields of research: studies on error detection, focussing on the recognition of errors, and studies on decision confidence, investigating processes related to beliefs regarding the likelihood of having made a correct choice. In most cases, low confidence implies a higher probability of having committed an error. It has been suggested that error detection and confidence judgements might even share similar underlying computations, whereby error detection arises from low confidence that a correct decision has been made (Boldt and Yeung, 2015; Yeung and Cohen, 2006; Yeung and Summerfield, 2014).

### 1.1 Neural correlates of metacognition

Neural correlates of metacognition have been studied by measuring event-related potentials (ERPs) of the human scalp electroencephalogram (EEG). The error negativity (N_e_) is a negative deflection peaking around 100 ms after an overt behavioural response at fronto-central electrodes and typically has larger amplitudes for errors than correct responses (N_c_ for correct responses; i.e., correct negativity; Falkenstein et al., 1991; Falkenstein et al., 2000; Vidal et al., 2003). The component is classically associated with conflict monitoring, assuming that it tracks conflict between the given response and continuously-accumulated post-decision evidence favouring the correct response (Falkenstein et al., 1991; Yeung et al., 2004). Moreover, it has been shown that the N_e_ amplitude scales with confidence, that is, it decreases from perceived errors to uncertain responses (guesses) to trials where the participant is confident about its correctness (Boldt and Yeung, 2015; Scheffers and Coles, 2000). The more posterior error positivity (P_e_; P_c_ for correct responses, i.e., correct positivity) with a maximum amplitude around 250 ms after a response, is considerably larger for detected compared to undetected errors and has therefore been associated with explicit error awareness (Endrass et al., 2012a; Nieuwenhuis et al., 2001). Notably, the P_e_ has also been found to increase in amplitude with decreasing confidence in perceptual decisions (Boldt and Yeung, 2015; Rausch et al., 2019).

Concerning the mechanisms underlying these two components, Di Gregorio et al. (2018) designed a sophisticated task to provide evidence that the P_e_, but not the N_e_, was present when it was evident for participants that an error had been made, but they did not know the correct answer. These findings suggest that the P_e_ does not require a representation of the correct response to emerge, but instead accumulates post-decisional error evidence from widely distributed neural sources (Di Gregorio et al., 2018; Murphy et al., 2015; Steinhauser and Yeung, 2010; Yeung and Summerfield, 2014). Thus, while both classical components of error processing, N_e_ and P_e_, have been shown to vary with reported confidence, the Pe appears to be more specifically associated with conscious metacognitive processes (Boldt and Yeung, 2015; Nieuwenhuis et al., 2001; Scheffers and Coles, 2000).

### 1.2 Metacognition and ageing

Metacognitive abilities in older age have been shown to vary across cognitive domains (Fitzgerald et al., 2017; Hertzog & Hultsch, 2000). For instance, while older adults tend to underestimate the prevalence of their decision errors in everyday life, metacognitive judgements of certain memory aspects (e.g., memory encoding) seem to be well preserved (Castel et al., 2016; Harty et al., 2013; Mecacci and Righi, 2006). Previous studies on decision making and metacognition yielded relatively consistent findings of a significant decline in error detection rate with higher age across multiple tasks (Harty et al., 2013; Rabbitt, 1990), even when task performance was comparable (Harty et al., 2017; Niessen et al., 2017; Wessel et al., 2018). In a large sample of healthy adults, Palmer et al. (2014) investigated decision confidence using a measure of metacognitive accuracy that takes task performance into account (Maniscalco & Lau, 2012). The authors found that age was not correlated with metacognitive abilities in a memory task, but that it was negatively correlated with metacognitive abilities in a perceptual discrimination task.

Effects of ageing on the neural correlates of metacognition have primarily been investigated in the field of error detection. Here, both the difference between N_e_ and N_c_ (Endrass, Schreiber, et al., 2012; Falkenstein et al., 2001; Schreiber et al., 2011), and the P_e/c_ amplitude (Clawson et al., 2017; Harty et al., 2017; Niessen et al., 2017) was smaller in older adults, while the decrease in P_e_, in particular, was linked to a lower error detection rate. Notably, the processing of the stimulus can also affect subsequent response-related processes, and variations with age in two ERPs (namely the N2 and the P300; Groom & Cragg, 2015; Polich, 2007) have been documented (Korsch et al., 2016; Larson et al., 2016; Lucci et al., 2013; Niessen et al., 2017). With the decline in behavioural performance reported above, this suggests an impaired error evidence accumulation process in older age, possibly due to limited cognitive resources (Harty et al., 2017; Niessen et al., 2017). Surprisingly, neither N_e/c_ nor P_e/c_ have been investigated using confidence ratings to assess age-related variations of metacognitive abilities. Some evidence from neuroimaging studies point to age-related structural differences in the neural basis of metacognition (Chua et al., 2009; Hoerold et al., 2013; Sim et al., 2020). However, a conclusive account that explains individual differences in metacognitive accuracy is still missing, for which the use of ERPs with high temporal resolution might be well-suited to provide valuable insights (Dully et al., 2018; Fleming and Dolan, 2012; Yeung and Summerfield, 2014).

### 1.3 The current study

This study aimed to investigate task performance and metacognition in older adults with a novel perceptual task to determine how generalizable the findings of decreased metacognitive accuracy in older age are (Palmer et al., 2014). For this, we used a colour-flanker task, in which participants had to identify the colour of a target stimulus that was flanked by two squares of the same or a different colour. We assessed decision accuracy, measured confidence using a four-point rating scale, and examined the impact of metacognitive accuracy on adaptations of subsequent behaviour (Desender et al., 2019a; Fleming et al., 2012; Ruitenberg et al., 2014). Furthermore, we investigated whether the amplitudes of N_e/c_ and P_e/c_, which are described as neural correlates of metacognition, track changes in decision confidence across the lifespan.

We hypothesised that metacognitive accuracy in our decision task would decrease with age (Niessen et al., 2017; Palmer et al., 2014). Independent of confidence, we expected an error-specific attenuation of ERP component amplitudes in older adults, which should result in a smaller difference between the neural responses related to errors and correct decisions (Endrass et al., 2012b; Larson et al., 2016). Independent of age, reported confidence was expected to show a positive association with the N_e/c_ and a negative association with the P_e/c_ amplitude (Boldt and Yeung, 2015; Scheffers and Coles, 2000). Based on findings from error detection studies showing an age-related decrease in the P_e_ amplitude of detected, but not undetected errors (Harty et al., 2017; Niessen et al., 2017), as well as reports linking the P_e_ to confidence (Boldt and Yeung, 2015), we expected a specific decrease in P_e_ amplitude for low confidence errors with increasing age.

## 2. Methods

### 2.1 Participants

We recruited 82 healthy adults with a broad age range from 20 to 81 years (49.8 ± 1.9 years [all results are indicated as mean ± standard error of the mean; *SEM*]; 35 female, 47 male). We aimed for an approximately uniform distribution of age and thus tested at least 10 participants per decade. Inclusion criteria were right-handedness according to the Edinburgh Handedness Inventory (EDI; Oldfield, 1971), fluency in German, (corrected-to-) normal visual acuity and no history of neurological or psychiatric diseases. Any signs of cognitive impairment (Mini-Mental-State Examination score lower than 24; MMSE; Folstein et al., 1975) or depression (Beck’s Depression Inventory score higher than 17; BDI; Hautzinger, 1991) led to the exclusion of participants (one participant was excluded). Additionally, we excluded four participants who had more than one third of invalid trials (e.g., responses were too slow to fall into the pre-defined response window for analysis, or they showed recording artefacts). Another four participants were excluded because of an error rate (ER) higher than the chance level of 75%. Finally, eight participants were excluded because of combinations of very low accuracy, a high number of invalid trials, the selective use of single response keys, and errors in the colour discrimination task (described below), which suggested a lack of understanding of the task or the use of heuristic response strategies instead of trial-by-trial decisions. After exclusions, the final sample consisted of 65 healthy adults (45.5 ± 2.0 years; 20 to 76 years; 26 female, 39 male).

The study was approved by the ethics committee of the German Psychological Society (DGPs) and conformed to the declaration of Helsinki. All participants gave written informed consent before participating in the experiment.

### 2.2 Experimental paradigm

The main experimental task consisted of a modified version of the Eriksen flanker task using coloured squares as stimuli and four response options (Eriksen & Eriksen, 1974; Maier & Steinhauser, 2017; Figure 1A). Participants were asked to respond as fast and accurately as possible to a centrally presented target by pressing a button with one of their index or middle fingers, mapped onto four designated target colours. In each trial, the target consisted of one of these target colours, and the flankers, located right and left to the target, consisted either of the same colour as the target (congruent condition), of another target colour (incongruent condition), or of one of three additional neutral colours, which were not mapped to any response (neutral condition [Maier et al., 2008]; see Figure 1B). Both the incongruent and the neutral condition were used to induce conflict as they provided information distinct from the target. We chose this version of the classical flanker paradigm in order to increase task difficulty and thereby maximise the number of errors without tapping into other cognitive processes that might be affected by ageing (e.g., spatial, lexical, or semantic cognition). The colour-finger mapping was fixed over the course of the experiment for each participant and counterbalanced across participants.

**Figure 1.**
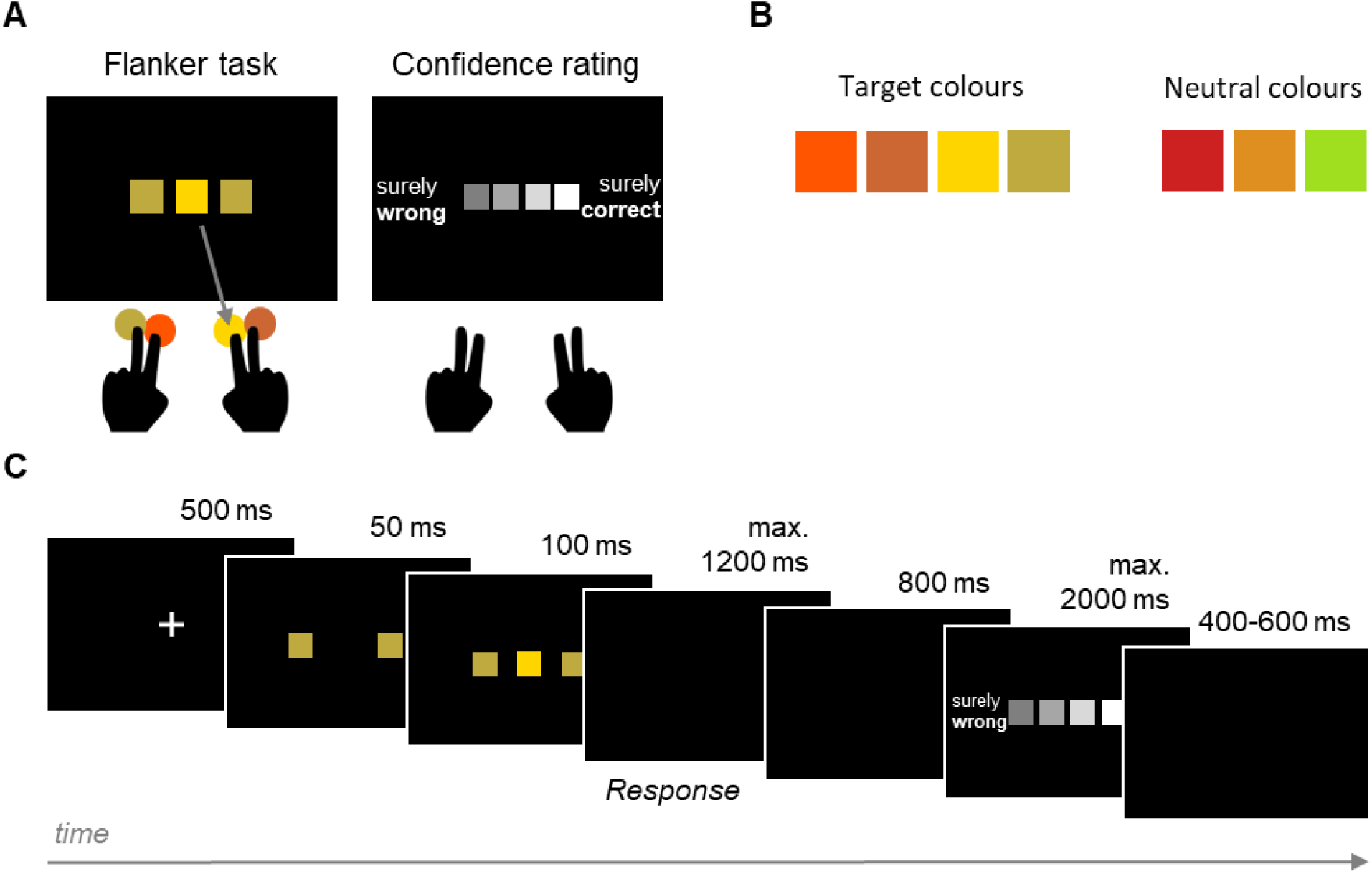
(A) The left panel shows an example of a trial in the flanker task, where one central target and two flankers were presented, and the participant had to press the finger that was assigned to the respective target colour (illustrated by the grey arrow). The confidence rating (right panel) consisted of four squares, and the ends of the scale were labelled with the German words for ‘surely wrong’ on the left and ‘surely correct’ on the right side. The fingers were mapped onto the four squares according to their spatial location. (B) Colours used in the flanker task. Flanker stimuli could consist of target or neutral colours, whereas target stimuli could only consist of one of the four target colours. (C) Sequence of one trial (here, incongruent). Each trial started with a fixation cross, followed by the presentation of the flankers, to which the target was added shortly after. Then, the screen turned black until a response was registered (maximum 1,200 ms), followed by another blank screen. If a response had been given, the rating scale appeared until a rating was given (maximum 2,000 ms). If no response had been given within the designated time window, the German words for ‘too slow’ were shown instead. The trial ended with another blank screen for a random intertrial interval.

Each trial started with a fixation cross for 500 ms. Then, flankers were presented for 50 ms before the target was added to the display for another 100 ms. Showing the (task-irrelevant) flankers before the target was expected to increase the induced conflict (Mattler, 2003). We used a response deadline of 1,200 ms because this timing provided a good balance between a desirable number of errors and feasibility for all participants. If no response was registered before this deadline, the trial was terminated and the feedback ‘zu langsam’ (German for ‘too slow’) was presented on the screen. If a response was given, a confidence rating scale appeared after a black screen of 800 ms. The delay was introduced to avoid that EEG activity related to the first response overlapped with the confidence assessment. Participants were asked to indicate their confidence in their decision on a four-point rating scale from ‘surely wrong’ to ‘surely correct’ using the same keys as for the initial response. The maximum time for the confidence judgment was 2,000 ms. Trials were separated by a jittered intertrial interval of 400 to 600 ms. The sequence of an experimental trial is depicted in Figure 1C.

### 2.3 Procedures

Prior to testing, participants were asked to provide demographic details and complete the handedness questionnaire. Afterwards, they completed a brief colour discrimination task (without EEG) to ensure that all participants were able to correctly discriminate the seven different colours used in the experimental paradigm (see Figure 1B). The discrimination task was followed by the EEG preparation and the main task. The neuropsychological tests were administered after the experiment. In addition, we assessed sustained attention span and processing speed using the d2-test (Brickenkamp, 2002), which have been shown to be positively associated with error processing abilities (Larson et al., 2011).

All stimuli in both tasks were presented on a black screen (LCD monitor, 60 Hz) in an electrically shielded and noise-insulated chamber with dimmed illumination, using Presentation software (Neurobehavioural Systems, version 14.5) for the colour discrimination task and uVariotest software (Version 1.978) for the main task. A chin rest ensured a viewing distance of 70 cm to the screen and minimised movements. To record participants’ responses, we used custom-made force-sensitive keys with a sampling rate of 1024 Hz (see Stahl et al., 2020).

The experiment started with a practice block of 18 trials in which participants received feedback about the accuracy of their response, which could be repeated if the participant considered it necessary. After that, two additional blocks with 72 trials without feedback and confidence assessments served as training blocks, allowing the participants to memorise the colour-finger mapping and to get accustomed to the response keys. Afterwards, another practice block introduced the confidence rating to ensure that participants understood and correctly applied the rating scale. The following main experiment consisted of five blocks with 72 trials each. Participants were allowed to take self-timed breaks after each block. The entire session lasted approximately three hours.

### 2.4 Electroencephalography recording and preprocessing

The EEG was recorded using 61 active electrodes (Acticap, Brain Products) aligned according to the international 10-20 system (Jasper, 1958). The electrodes were online referenced against the posterior Iz electrode close to the inion. Horizontal eye movements were measured using two electrodes at the outer canthi of the eyes (horizontal electrooculogram [EOG]), and another electrode underneath the left eye measured vertical movements (vertical EOG). The EEG signal was recorded continuously at a sampling rate of 500 Hz using a digital BrainAmp DC amplifier (Brain Products). Data were filtered between 0.1 Hz and 70 Hz, and a notch filter of 50 Hz was applied to remove line noise.

EEG data were preprocessed following a standardised pipeline using the MATLAB-based toolboxes EEGLAB and ERPLAB (Delorme and Makeig, 2004; Lopez-Calderon and Luck, 2014). The signal was segmented from −150 to 2,000 ms relative to target stimulus presentation (note that the flankers were presented at −50 ms). Epochs were visually inspected for artefacts and noisy electrodes. Epochs with artefacts were removed and identified noisy channels were interpolated using spherical spline interpolation. To identify and remove eyeblinks, we ran an Independent Component Analysis (ICA) using the infomax algorithm implemented in EEGLAB and afterwards baseline-corrected the epochs using the period of −150 ms to −50 ms to avoid influences of early perceptual processes related to the flanker presentation. Next, data were locked to the response, epoched from −150 ms to 800 ms relative to response onset and baseline-corrected using the 100 ms before the response. The additional analysis of conflict-related stimulus-locked ERPs can be found in the supplementary material S3. Remaining artefacts exceeding ± 150 μV were removed (Niessen et al., 2017), and a current source density (CSD) analysis was conducted using the CSD toolbox (Kayser and Tenke, 2006) allowing for better spatial isolation of ERP components and for obtaining a reference-independent measure (Perrin et al., 1989).

### 2.5 Behavioural data analysis

Trials with invalid responses (i.e., responses that were too slow) or recording artefacts, as well as responses faster than 200 ms were excluded from further analysis. The error rate (ER) was calculated as the proportion of errors relative to valid responses. Response time (RT) was defined as the time between stimulus onset and the initial crossing of the force threshold (40 cN) by any of the response keys. For pre-defined conditions of interest (see below), individual median RTs were computed, and the means across participants were entered into group-level statistical analyses.

To inspect how the confidence scale was used across participants, raw distributions of confidence ratings within all incorrect and correct responses were extracted. We computed Friedman ANOVAs for the percentage of each of each rating level for errors and correct responses with the factor confidence (4 levels). This analysis revealed that only a limited number of trials was available for the two middle confidence rating levels (‘maybe wrong’, ‘maybe correct’), and we therefore collapsed those to create one category for all further analyses representing ‘unsure’ responses, i.e., confidence ratings expressing uncertainty.

For the analysis of metacognitive accuracy, we computed the Phi (Φ) correlation coefficient, which is a simple trial-wise correlation between task accuracy and reported confidence. It describes the extent to which the distributions of confidence ratings for correct and incorrect trials differ (Fleming and Lau, 2014; Kornell et al., 2007; Nelson, 1984). Phi was calculated by correlating accuracy, coded as 0 (error) and 1 (correct response), and confidence (that the given response was correct), coded as 1 (‘surely wrong’), 2 (‘unsure’), and 3 (‘surely correct’), for each participant. This provided us with one measure of metacognitive ability per participant that comprises both the accuracy and the confidence rating of each trial (e.g., Phi = 1 means that correct trials were successfully identified as such without uncertainty; while a Phi = 0 means that all errors were rated as ‘surely correct’, or all correct trials were rated as ‘surely incorrect’).

To assess the impact of accuracy and confidence on trial *n* on adaptations of behavioural responses, we computed a measure of response caution on trial *n*+1 by multiplying the individual percentage correct and median RT (Desender et al., 2019a). Response caution captures the trade-off between speed and accuracy in a decision, with higher values indicating a more cautious response strategy that is characterised by slower, and at the same time, more accurate responses. For this analysis, only to pairs of consecutive valid trials were included. Response caution was computed separately relative to a) initial trial accuracy (error, correct), and b) each confidence category (‘surely wrong’, ‘unsure’, ‘surely correct’) of the initial trial.

Age-related effects on the d2-test score, the error rate, and Phi were computed using Pearson correlations between these behavioural variables and age. To rule out that metacognitive accuracy was confounded by task performance or attention and processing speed, we computed partial correlations between Phi and age, controlling for the individual error rate and d2-test score, respectively.

For the analysis of performance and confidence across the lifespan, we performed a series of one-way repeated measures ANCOVAs with age as the between-subject covariate of interest. The dependent variables were error rate, RT, and response caution, and either accuracy (error, correct), or (pooled) confidence (3 levels) served as the within-subject factor.

Significant main effects of accuracy or confidence were followed up by planned paired-samples *t*-tests comparing the dependent variable between all levels. Significant interactions with age were followed up by Pearson correlations, separately for each level of a given within-subject factor. We used these follow-up tests because our main interest was in the differential relations between accuracy, confidence, and behaviour across the lifespan (rather than between the levels).

Post-hoc tests were computed using Bonferroni corrected *p*-values. If the assumption of sphericity was violated, degrees of freedom were corrected according to Greenhouse-Geisser, and adjusted *p*-values are reported. Effect sizes are reported as partial eta squared (η^2^_p_), Cohen’s d, or Pearson’s correlation coefficient r, respectively. Analyses were run in MATLAB R2019a (The Mathworks, Inc.) and IBM SPSS Statistics for Windows (Version 25.0).

### 2.6 Electroencephalographic data analysis

Data were response-locked and averaged for each participant, separately for the following categories: a) errors and correct responses (across confidence levels), b) errors (‘low confidence’ and ‘higher confidence’), and c) correct (‘high confidence’ and ‘lower confidence’). Error trials rated as ‘surely wrong’ were termed ‘low confidence’; error trials which received any higher confidence rating were termed ‘higher confidence’ (and vice versa for correct trials). The reason for aggregating the trials in this way rather than using three levels as for the behavioural analyses was that errors were most often associated with a ‘surely wrong’ and correct trials with a ‘surely correct’ rating, causing trial numbers for the other three rating categories to be too low to be included as separate categories of interest (note that because of data cleaning, typically less trials are available for the analysis of ERP data).

The N_e/c_ local peak amplitude was extracted from the response-locked data from the interval 0 to 150 ms following the response at FCz, and the P_e/c_ local peak amplitude was extracted from the interval 150 to 350 ms at Cz based on conventions and visual inspection of the local maxima of the grand-average scalp topographies (Falkenstein et al., 2000; Siswandari et al., 2019).

Only data sets with a minimum of six trials in each condition of the respective comparison were used for statistical analyses (Pontifex et al., 2010). To maximise the number of data-sets available for each analysis of interest, three subgroups of participants were used: Subgroup 1 included 63 participants and was used to examine the effect of accuracy and age on the ERP amplitudes for pooled trials. In addition, we ran focussed control analyses on two smaller subgroups of participants who had a sufficient number of trials to investigate modulations by confidence across the lifespan separately for errors (Subgroup 2, *N* = 44) and correct responses (Subgroup 3, *N* = 51). For the latter analyses, we ensured that participants’ age was still well-distributed across all decades within the subgroups, and that age similarly affected the main behavioural parameters as in the full sample (see supplementary material S1).

For statistical analyses of ERP peak amplitudes, we also employed one-way repeated measures ANCOVAs with age as the covariate of interest. For Subgroup 1, the dependent variables were the CSD-transformed N_e_ and N_c_ as well as P_e_ and P_c_, amplitudes with accuracy (error, correct) as the within-subject factor (see supplementary material S2 for the analysis with the within-subject factor confidence). For Subgroup 2 (including only errors), the dependent variables were N_e_ and P_e_, and the within-subject factor was confidence (‘surely wrong’, ‘higher confidence’), and lastly, for Subgroup 3 (including only correct responses), the dependent variables were N_c_ and P_c_, and the within-subject factor was again confidence; however, with different levels (‘surely correct’, ‘lower confidence’).

## 3. Results

Only significant effects in the ANCOVAs and relevant follow-up tests are reported in this section. For results of all tests, please refer to supplementary Tables S1-S4. Note that all computed ANCOVAs included age as the covariate of interest.

### 3.1 Behavioural results

#### 3.1.1 Attention

The average score for sustained attention and processing speed as assesses by the d2-test was 178.5 ± 5.6 (*M* ± *SEM*) and showed the typical decline for older adults, as shown by a significant correlation between test scores and age [*r*(63) = −.553, *p* < .001].

#### 3.1.2 Distribution of confidence ratings

In a first step, we were interested in how the confidence ratings were distributed across the four confidence levels across the lifespan (Figure 2). For this, we ran two Friedman ANOVAs for dependent measures for the percentages for each rating category, separately for errors and correct responses.

**Figure 2.**
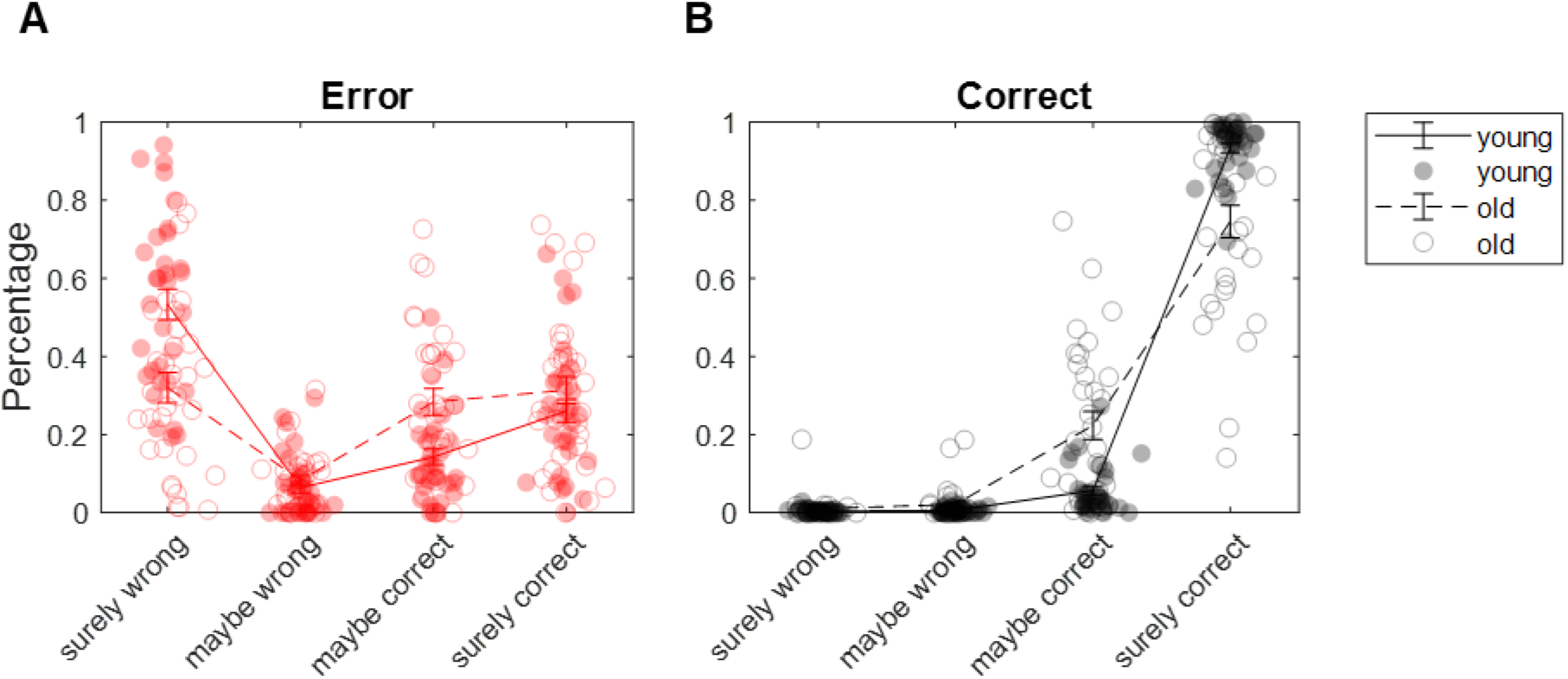
Distribution of confidence ratings for errors (red; A) and correct responses (black; B). Errors were most often rated as ‘surely wrong’, and correct responses as ‘surely correct’. Dots represent the individual proportion of the particular confidence response amongst all errors or correct responses, respectively. A median split by age (*Mdn* = 46) was conducted for illustration purposes. Filled dots and solid lines represent younger adults, empty circles and dashed lines represent older adults. With increasing age, participants used the ‘surely correct/wrong’ ratings less, but more the middle of the confidence scale.

The ANOVA for errors showed that the percentages differed between confidence levels [*Χ^2^*(3) = 78.029, *p* < .001]. On average, most errors were rated as ‘surely wrong’ (42.8 %) and the smallest proportion of errors as ‘maybe wrong’ (7.3%). Interestingly, follow-up correlation analyses between age and percentage for each rating category showed that the proportion of ‘maybe correct’ ratings (21.0%) was increased with higher age [*r*(63) = .450, *p* < .001], whereas the ratio of ‘surely wrong’ ratings was decreased [*r*(63) = −.543, *p* < .001; all other correlations *ns*; Figure 2A].

For correct responses (Figure 2B), the ANOVA also revealed a main effect of confidence [*Χ^2^*(3) = 167.472, *p* < .001]. Correct responses were most often rated as ‘surely correct’ (84.2 %) and least often as ‘surely wrong’ (0.7 %). Again, correlation analyses between age and percentage within each confidence category showed that the proportion of ‘maybe correct’ ratings (13.8%) was increased with higher age [*r*(63) = .530, *p* < .001], and was decreased for ‘surely correct’ ratings [*r*(63) = −.532, *p* < .001; all other correlations *ns*].

As mentioned above, to ensure a sufficient number of trials for each level of confidence for each participant, we combined ‘maybe wrong’ and ‘maybe correct’ ratings into one category representing ‘unsure’ responses. Thus, for all following behavioural analyses including the factor confidence, the reported analyses use three confidence levels.

#### 3.1.3 Error rate (ER)

The average error rate was 15.6 ± 1.6% and significantly increased with higher age [*r*(63) = .594, *p* < .001]. The ANCOVA for error rate with the within-subject factor confidence revealed a main effect of confidence [*F*(2,124) = 97.426, *p* < .001, η^2^_p_ = .611]. The error rate decreased across confidence levels from 92.1 ± 1.4% on trials rated as ‘surely wrong’ to 34.7 ± 2.6% on trials rated as ‘unsure’ and 8.1 ± 1.4% on trials rated as ‘surely correct’. This showed that, on average, participants’ confidence reflected their performance reasonably well (which further supports the notion that the current study’s confidence scale was a meaningful assessment tool). Furthermore, the ANCOVA revealed a significant interaction between confidence and age [*F*(2,124) = 5.264, *p* = .009, η^2^_p_ = .078]. In subsequent correlation analyses between error rate and age for each level of confidence, a higher error rate with older age was only found for the ‘surely correct’ confidence level [*r*(63) = .568, *p* < .001; both other correlations *ns*].

#### 3.1.4 Response time (RT)

An ANCOVA for mean RT with the within-subject factor accuracy showed the expected slowing with age [*F*(1,63) = 18.164, *p* < .001, η^2^_p_ = .224; correlation between age and RT: *r*(63) = .534, *p* < .001] and a significant main effect of accuracy [*F*(1,63) = 4.188, *p* = .045, η^2^_p_ = .062]. A follow-up *t-*test revealed that errors (733.4 ± 13.9 ms) were on average slower than correct responses [709.1 ± 11.5 ms*; t*(64) = 2.937, *p* = .005, *d* = 0.364]. The interaction between accuracy and age was not significant.

The age-related slowing was similarly present as a main effect of age in the ANCOVA with the within-subject factor confidence [*F*(1,57) = 19.159, *p* < .001, η^2^_p_ = .252; Figure 4A]. Moreover, the ANCOVA revealed a main effect of confidence [*F*(2,114) = 13.132, *p* < .001, η^2^_p_ = .187] and a significant interaction between confidence and age [*F*(2,114) = 3.793, *p =* .030, η^2^_p_ = .062]. Planned follow-up *t*-tests showed that trials associated with the ‘unsure’ confidence level (805.4 ± 15.0 ms) were significantly slower than trials rated as ‘surely correct’ [700.0 ± 11.7 ms; *t*(58) = 9.198, *p* < .001, *d* = 1.198] or ‘surely wrong’ [725.2 ± 16.9 ms; *t*(58) = −5.236, *p* < .001, *d* = 0.682]. Furthermore, trials with the extreme ratings [‘surely wrong’: *r*(57) = .502, *p* < .001; ‘surely correct’: *r*(57) = .567, *p* < .001], but not with ‘unsure’ ratings, were significantly slower with older age.

In short, RT was associated with confidence, such that high certainty (i.e., ‘surely correct/wrong’) was associated with the fastest responses, and this difference decreased with higher age.

#### 3.1.5 Confidence

A repeated measures ANCOVA examining mean confidence ratings (coded from 1 to 3, i.e., ‘surely wrong’, ‘unsure’, and ‘surely correct’) across all trials revealed a main effect of accuracy [i.e., error vs. correct trials; *F*(1,63) = 164.008, *p* < .001, η^2^_p_ = .722] and a significant interaction between accuracy and the covariate age [*F*(1,63) = 37.433, *p* < .001, η^2^_p_ = .373]. The average confidence rating was lower for errors (1.861 ± 0.047) compared to correct responses (2.836 ± 0.026), as confirmed in a follow-up *t*-test [*t*(64) = −16.774, *p* < .001, *d* = 2.081]. Separate follow-up correlation analyses between confidence and age for each accuracy level separately (error, correct) showed that the mean confidence for errors was increased with higher age [*r*(63) = .471, *p* < .001], while the mean confidence was decreased with higher age for correct responses [*r*(63) = −.523, *p* < .001; Figure 3B].

**Figure 3.**
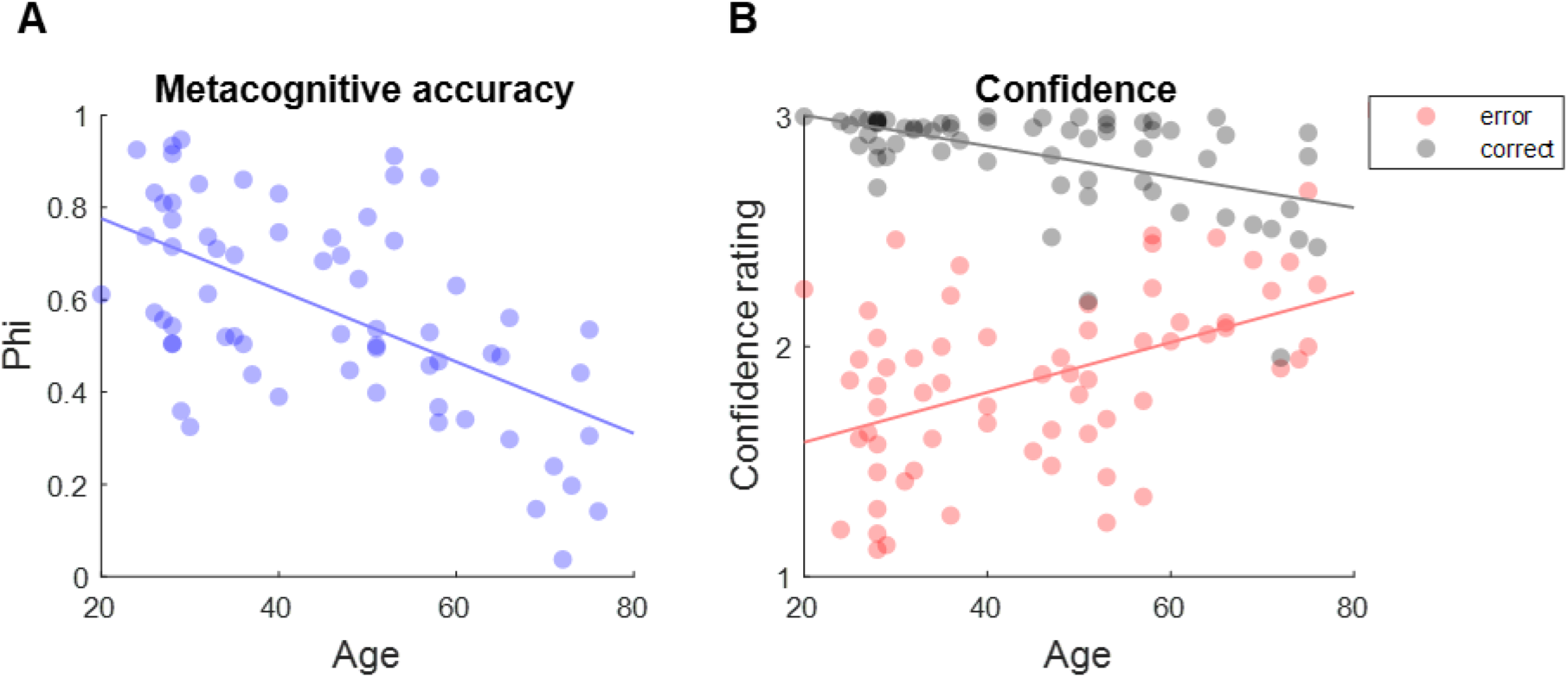
Metacognition across the lifespan. (A) Metacognitive accuracy (Phi) decreased with age. (B) Mean confidence ratings of errors correlated significantly positively with age, while mean confidence ratings of correct trials correlated significantly negatively. Dots represent means of individual participants.

**Figure 4.**
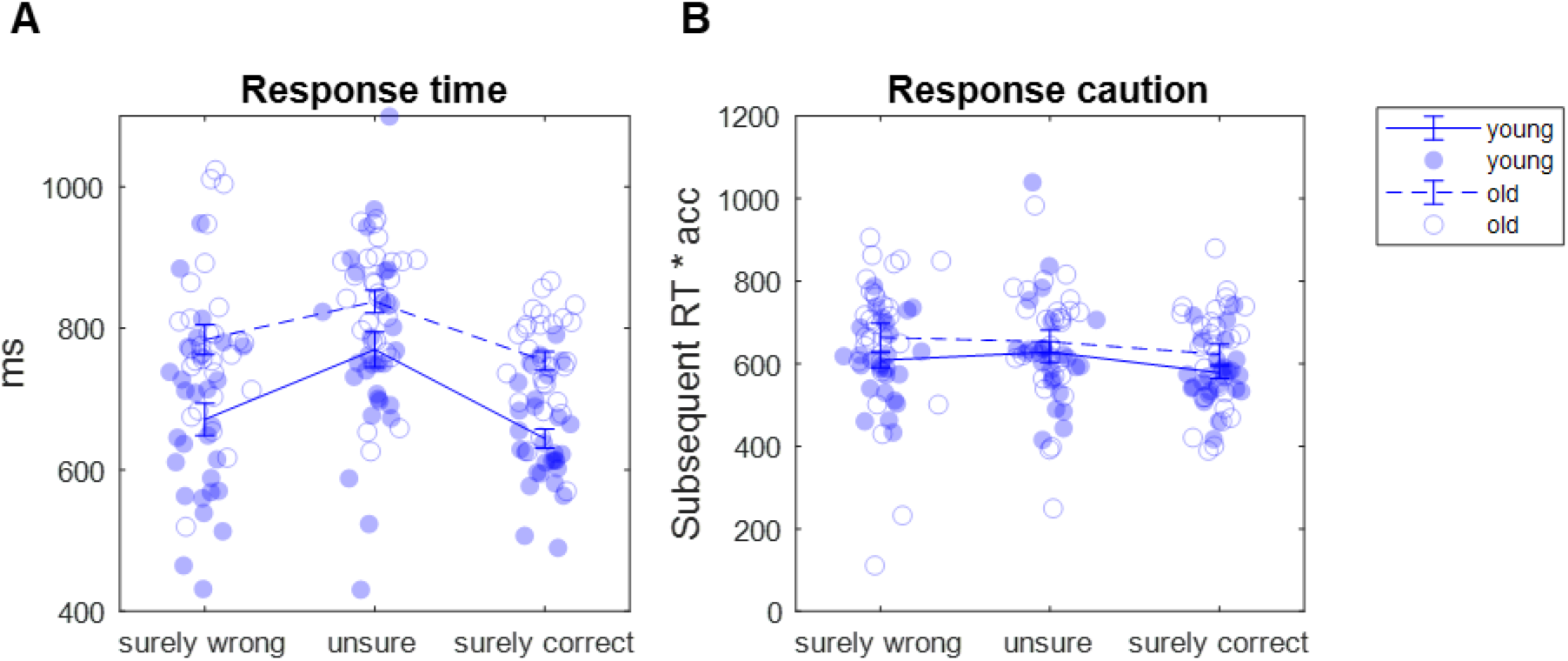
Modulation of response time (RT; A) and response caution (B) by confidence. Participants were split into a younger and an older group relative to the median (*Mdn* = 46) for illustration purposes. Filled dots and solid lines represent younger adults, empty circles and dashed lines represent older adults. (A) Trials rated as ‘unsure’ showed slower RTs than trials associated with a ‘surely’ rating, and this difference was smaller with increasing age. (B) Adaptation of response caution depending on previous trial confidence rating. Response caution was computed as the product of individual average accuracy and RT of subsequent trials.

#### 3.1.6 Metacognitive accuracy (Phi)

Phi had a mean of 0.579 ± 0.027 across the entire sample and correlated negatively with age, indicating a decrease of metacognitive accuracy with age [*r*(63) = −.582, *p* < .001; Figure 3A]. Moreover, we computed a partial correlation between Phi and age while controlling for error rate, which still showed a decrease of Phi with age [*r*(60) = −.343, *p* = .005], suggesting that changes in metacognitive accuracy with age were not due to individual differences in task performance. Similarly, a partial correlation controlling for the individual d2-test scores (which provide a task-independent measure of attention) suggested that the decrease in Phi with age was also independent of an age-related reduction in attentional capacity [*r*(60) = −.487, *p* < .001].

#### 3.1.7 Behavioural adaptation

To investigate the effect of accuracy and confidence in a given trial on the behaviour in the following trial, we computed response caution as the product of individual percentage correct and median RT in the subsequent trial. The ANCOVA with the within-subject factor accuracy (referring to the previous trial) revealed a main effect of accuracy [*F*(1,63) = 6.929, *p* = .011, η^2^_p_ = .099] but no main effect of age, nor a significant interaction. The main effect was confirmed in a follow-up *t*-test between errors and correct responses [*t*(64) = 2.950, *p* = .004, *d* = 0.357]. These findings indicate that participants were on average more cautious after errors than after correct responses, and this effect was independent of age.

Next, we examined whether the response caution in the subsequent trial could also be modulated by confidence. As shown above, confidence and accuracy are strongly correlated; however, a significant modulation by confidence could also indicate that this internal confidence signal drives behavioural adaptations (Figure 4B). The ANCOVA with the within-subject factor confidence (referring to the previous trial) revealed a trend for a main effect of confidence [*F*(2,110) = 2.897, *p* = .059, η^2^_p_ = .050], but again, no main effect of age, nor a significant interaction. Follow-up *t*-tests between the confidence levels showed that the response caution after trials rated as ‘surely correct’ was significantly lower compared to trials rated as ‘unsure’ [*t*(56) = 3.448, *p* = .001, *d* = 0.457] or as ‘surely wrong’ [*t*(56) = 3.066, *p* = .003, *d* = 0.406].

To summarise the effects of ageing on behaviour, we found the expected age-related general increase in error rates and response times, accompanied by a decrease in metacognitive ability, which was mainly reflected in reduced use of confidence ratings at the extreme ends of the scale but more indications of being unsure. Response caution, on the other hand, was not affected by ageing. Caution increased after errors compared to correct responses, and notably, tended to be specifically modulated by previous trial confidence. With higher confidence, the response caution in the subsequent trial decreased.

### 3.2 Electrophysiological results

#### 3.2.1 N_e/c_ amplitudes

The peak amplitude of the N_e/c_ was significantly larger for errors compared to correct responses, as reflected in a main effect of accuracy in the ANCOVA with the within-subject factor accuracy and the covariate of interest age [*F*(1,61) = 15.209, *p* < .001, η^2^_p_ = .200; *t*(62) = −4.544, *p* < .001, *d* = 0.572; Figure 5A & B]. A main effect of age revealed smaller amplitudes with older age [*F*(1,61) = 5.999, *p* = .029, η^2^_p_ = .076], and an interaction between accuracy and age showed that the age effect differed between errors and correct responses [*F*(1,61) = 5.999, *p* = .017, η^2^_p_ = .090]. Follow-up correlation analyses between the N_e_ and N_c_ amplitude and age, separately for errors and correct responses, confirmed that the N_e_ [*r*(61) = .326, *p* = .009], but not the N_c_ amplitude was reduced with higher age.

**Figure 5.**
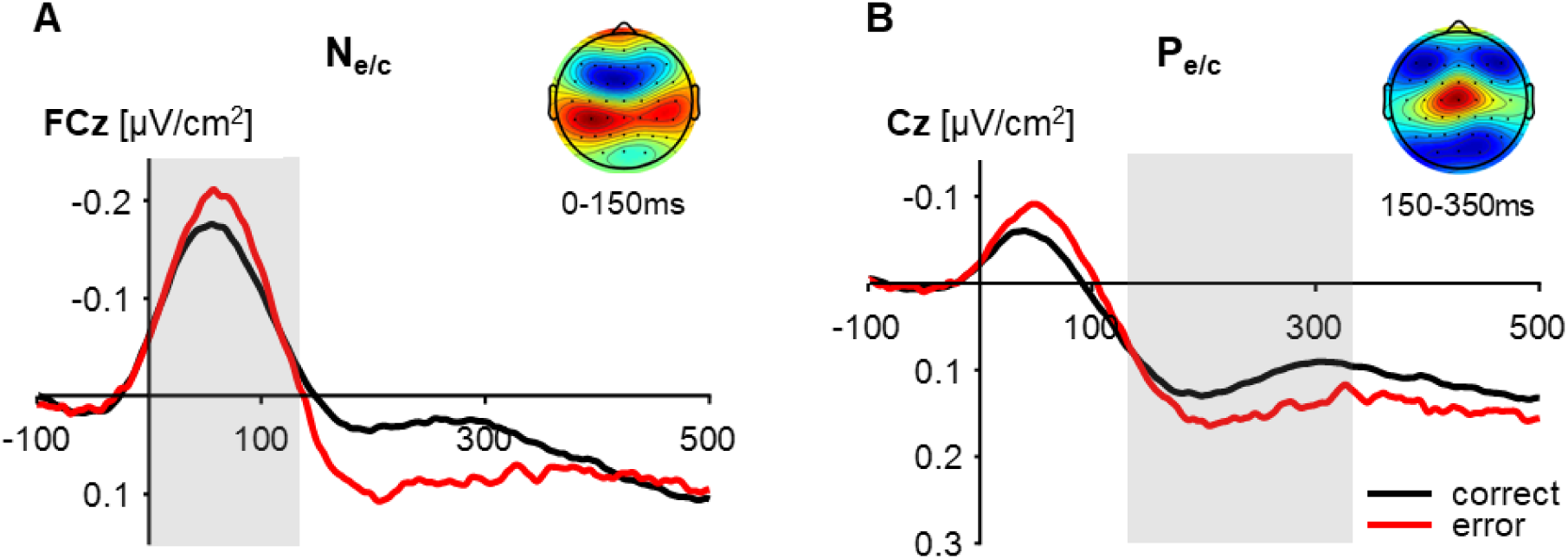
Response-locked event-related potentials for errors and correct responses and topographical maps of errors (*N* = 63; A, B) after current source density transformation. (A) N_e/c_ is computed at electrode FCz and (B) P_e/c_ at electrode Cz. Errors are shown in red, correct trials in black. Scalp topographies depict the mean activity for all error trials averaged across the time windows for the N_e_ (0-150 ms) and the P_e_ (150-350 ms). Grey squares indicate time windows for the analysis of peak amplitudes for the respective components.

For the analysis of confidence, we ran separate ANCOVAs for errors and correct responses on N_e_ and N_c_ amplitude, respectively. Confidence served as the within-subject factor, but was collapsed into two levels (error: ‘surely wrong’, ‘higher confidence’; correct: ‘surely correct’, ‘lower confidence’; see Methods). The ANCOVA for errors showed a main effect of age [*F*(1,42) = 10.787, *p* = .002, η^2^_p_ = .204], but the main effect of confidence failed to reach significance [*F*(1,42) = 3.461, *p* = .070, η^2^_p_ = .076]. No significant interaction was found (Figure 6A). Follow-up *t*-tests showed that N_e_ amplitudes were indeed larger for errors of low confidence compared to higher confidence [*t*(43) = −2.309, *p* = .026, *d* = 0.348].

**Figure 6.**
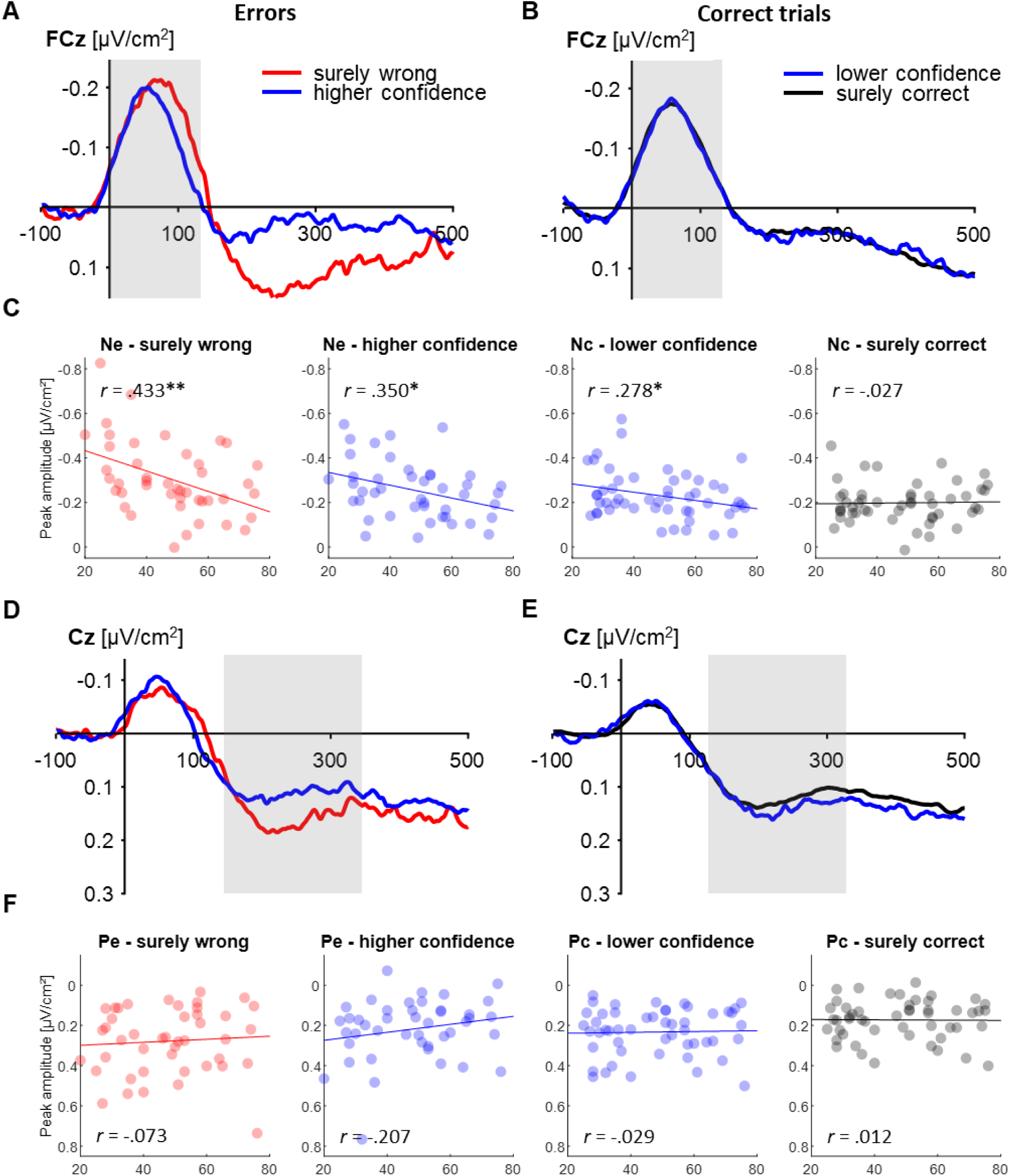
Response-locked event-related potentials, separately for errors and correct responses after current source density transformation. Errors are shown in the left panel, correct trials in the right panel. For errors (*N* = 44), data are binned into trials rated as ‘surely wrong’ (red) and trials rated with any higher confidence (blue). The N_e_ (A) and N_c_ (B) are shown at electrode FCz and did not differ between confidence levels, and P_e_ (D) and P_c_ (E) are shown at electrode Cz and were both increased for the trials associated with lower confidence, respectively. (C) and (F) illustrate the correlations between age and the ERP components, separately for errors, correct trials, and confidence levels. The amplitudes of the N_e_ (both confidence levels) and the N_c_ (only ‘lower confidence’) decreased with age, while there was no significant correlation between age and any of the P_e/c_ amplitudes. Dots represent individual average peak amplitudes. \**p* < .05. \*\**p* < .01. \*\*\**p* < .001

The ANCOVA for correct responses with the within-subject factor confidence yielded a significant effect of confidence [*F*(1,49) = 12.624, *p* = .001, η^2^_p_ = .205]. The N_c_ was not modulated by age, but we found a significant interaction between confidence and age [*F*(1,49) = 7.746, *p* = .008, η^2^_p_ = .136; Figure 6B]. The main effect was driven by smaller N_c_ amplitudes after correct responses rated as ‘surely correct’ compared to those with lower confidence, as reflected in a significant follow-up *t*-test [*t*(50) = −2.699, *p* = .009, *d* = 0.438]. Follow-up correlation analyses pointed to an age-related decrease in N_c_ amplitude on correct trials with lower confidence ratings [*r*(49) = .278, *p* = .048], but not when they were rated as ‘surely correct’ (Figure 6C).

#### 3.2.2 P_e/c_ amplitudes

The ANCOVA for the P_e/c_ with the within-subject factor accuracy revealed a significant main effect for accuracy with larger amplitudes for errors compared to correct responses [*F*(1,61) = 23.886, *p* < .001, η^2^_p_ = .281; *t*(62) = 4.507, *p* < .001,*d* = 0.568; Figure 5C & D]. There was no main effect for age, but a significant interaction between confidence and age [*F*(1,61) = 11.836, *p* = .001, η^2^_p_ = .163]. Follow-up correlation analyses for errors and correct responses found no significant correlation between the amplitude and age for correct responses; however, the age-related decrease in amplitude for errors marginally missed the significance threshold [*r*(61) = −.237, *p =* .061].

Next, responses were again split by their accuracy, and separate ANCOVAs were conducted with the within-subject factor confidence. The ANCOVA for errors did not yield any significant effects on the P_e_ amplitude (Figure 6D). However, due to previous evidence of a strong relation between P_e/c_ amplitude and error detection or confidence ratings (Boldt and Yeung, 2015; Nieuwenhuis et al., 2001), we performed an exploratory *t*-test comparing the P_e_ amplitudes between the two confidence levels. Indeed, we found a significant difference between ‘surely wrong’ and ‘higher confidence’ errors [*t*(43) = 2.157, *p* = .037, *d* = 0.325], suggesting that the P_e_ was only modulated by confidence, if assessed independent of age.

The ANCOVA for correct trials, in contrast, revealed a main effect of confidence [*F*(1,49) = 5.065, *p* = .029, η^2^_p_ = .094], and follow-up *t*-tests showed that the peak amplitude was on average higher for correct responses with lower confidence compared to those rated as ‘surely correct’ [*t*(50) = 5.838, *p* < .001, *d* = 0.818; Figure 6E]. Again, there was no effect of age on P_c_ amplitude, neither as a main effect, nor as an interaction.

## 4. Discussion

We conducted a complex four-choice flanker task with adult participants covering an age range from 20 to 76 years, allowing us to investigate confidence and metacognitive accuracy as well as neural indices thereof across the lifespan. We found that error rates and response times (RT) increased with age. Metacognitive accuracy, quantified as Phi, gradually decreased across the lifespan and was characterised by differential use of confidence ratings. In contrast, we did not find differences between younger and older adults in the ability to adapt behaviour in accordance with reported confidence. As expected, the N_e/c_ and P_e/c_ amplitudes declined with higher confidence. While the N_e/c_ amplitude was smaller with older age whenever participants were not entirely certain about their response accuracy, the variation in the P_e/c_ amplitude with reported confidence was surprisingly not affected by ageing. In the following, we will first discuss potential processes underlying age-related differences in metacognitive accuracy and their relation to task performance and confidence, before comparing the pattern we observed at the behavioural level to the patterns we observed in the ERPs. Finally, we argue that older adults’ preserved ability to adapt their behaviour to their perceived confidence could be related to the P_e/c_ amplitude.

### 4.1 Differential use of confidence scale as a marker of age-related metacognitive decline

In the present study, metacognitive accuracy (Phi) was reduced with increasing age. This is consistent with the findings of Palmer et al. (2014) who used a metacognitive efficiency measure, which further considered the individual performance in their perceptual discrimination task. As this measure was not directly applicable in our four-choice flanker task, we confirmed (by calculating partial correlations taking into account the error rate and the d2-test score) that the observed decline in metacognitive accuracy was not merely a reflection of general age-related performance or attention deficits (d2-test; see also Larson & Clayson, 2011). Our results, therefore, show that Palmer et al.’s (2014) findings also hold for a more complex, speeded decision task, which was not based on stimulus ambiguity.

The question remains as to how the age-related differences in confidence emerge. Given the nature of Phi, a smaller value could either indicate more ‘misclassifications’, or a general response tendency towards the middle (i.e., rating all correct responses as ‘maybe correct’ will result in a lower Phi value than rating the same number of correct responses as ‘surely correct’). Indeed, we observed that older adults used the extreme ends of the confidence scale considerably less often than younger adults.

For errors, this pattern resulted in a higher mean confidence with age. This disproportional rise in reported confidence has similarly been shown in error detection studies, indicated by a lower error detection rate in older adults (Harty et al., 2017, 2013; Niessen et al., 2017).

For correct decisions, in contrast to previous studies that reported an *over*-confidence in older age (Dodson et al., 2007; Hansson et al., 2008; Ross et al., 2012), we observed *lower* mean confidence due to the tendency of the older adults to use the middle of the confidence scale more often. These findings emphasise the actual difficulties of older adults in evaluating their performance and establishing confidence. In our study, the (relatively) high task difficulty might be a reason for these difficulties. Stahl et al. (2020) showed that their slow errors, which were associated with a weak stimulus-response representation (i.e., due to a weak memory), were associated with lower confidence than fast, impulsive responses. Thus, assuming that the present task posed higher demands on the older adults (as indicated, for instance, by higher error rates), their impaired metacognitive evaluation might partly be related to more frequent memory-related errors, which appear to be more challenging to assess consciously (Maier and Steinhauser, 2017; Stahl et al., 2020).

### 4.2 Neural correlate of confidence is stable across age

The P_e/c_ is an established marker of metacognition, reflecting variations in subjective error awareness and decision confidence (Boldt and Yeung, 2015; Nieuwenhuis et al., 2001). In the present study, the P_e/c_ showed the well-known accuracy effect of larger amplitudes for errors than correct responses. Moreover, we could replicate prior findings of the P_e/c_ increasing with decreasing confidence, - for the first time - for a very broad age range (Boldt and Yeung, 2015; Rausch et al., 2019). The modulation of the P_e_ of errors by confidence also replicates findings from error detection studies showing increased P_e_ amplitudes for detected compared to undetected errors (Endrass et al., 2012a; Nieuwenhuis et al., 2001).

The main interest of our study was to investigate the modulation of the P_e/c_ by metacognition in the context of healthy ageing. Remarkably, the P_e/c_ amplitude did not show an overall reduction with age, nor a differential modulation by confidence across the lifespan, suggesting that the accumulation of error evidence was well preserved in older age. This is contrary to the error detection literature (Harty et al., 2017; Niessen et al., 2017). Since these studies did not assess confidence on multiple levels, participants did not have the chance to express uncertainty. Assuming more ‘unsure’ cases with older age, their observed age-related decrease in P_e_ amplitude for detected errors might thus be confounded, as higher uncertainty was generally associated with reduced P_e_ amplitudes (Boldt and Yeung, 2015). Following this logic, it is also possible to explain the lack of a significant age-related modulation of the P_e/c_ amplitude in the present study: If older adults’ internal threshold for rating an error as ‘surely wrong’ was generally raised, the errors that *were* rated as ‘surely wrong’ should be trials with particularly high P_e_ amplitudes, as they were absolutely sure of having committed an error. As a result, a putative age-related decrease in the P_e_ amplitude of low confidence errors could be masked in our data, because the same reported rating levels might reflect a different sense of confidence for younger and older adults. Thus, the current pattern of results suggests that the P_e/c_ amplitude does *not* serve as a direct index of metacognitive accuracy across participants, but rather reflects the degree of confidence, irrespective of objective performance (Di Gregorio et al., 2018; Larson and Clayson, 2011; Pouget et al., 2016; Stahl et al., 2020).

### 4.3 Impaired neural processing of conflict modulates metacognitive decline

The marked behavioural decline in older adults’ metacognitive accuracy was not mirrored in age-related variations of the P_e/c_ amplitude, but rather in a differential modulation of the N_e/c_ across the lifespan. The control analyses revealed that the N_e/c_ amplitude was also affected by the interaction between confidence and age. With older age, the N_e/c_ declined for all trial types in which some degree of conflict was perceived. In other words, only the N_c_ of correct trials that were rated as ‘surely correct’ showed no amplitude variations across the lifespan. As the N_e/c_ is sensitive to conflict between the given and the actual correct response, older adults seemed to having had difficulties internally representing the correct response in any conflicting situation (Yeung et al., 2004).

We suggest that the reduced N_e/c_ amplitude of perceived errors with higher age could be related to the observed decrease in metacognitive accuracy in our flanker task. If older adults had difficulties forming an accurate internal representation of the correct response, this information was necessarily missing for the metacognitive evaluation. Thus, the impaired neural integration of conflict detection and confidence could have led to the observed behavioural difficulties matching confidence ratings and objective accuracy.

### 4.4 Adults of all ages base future behaviour on subjective confidence

Ultimately, proper metacognitive evaluation should improve behaviour. Interestingly, response caution was not only enhanced after errors, but we also found some (limited) evidence that it was modulated by the reported confidence in the preceding trial. Given that the participants did not receive any external feedback about the accuracy of their response (as it is often the case in real-life decisions), it seems plausible that they used their best available estimate, i.e., the subjective sense of confidence, to regulate subsequent behaviour (Desender et al., 2019a). Specifically, medium and low confidence about a choice was associated with higher response caution in the subsequent trial. Possibly, participants sought more evidence before committing to their next decision, leading to slower but more accurate responses (Desender et al., 2019a, 2019b).

Translating our findings to error detection studies, the increase in response caution with lower previous trial confidence converges with findings of error detection studies reporting increased slowing (i.e., a sign of behavioural adaptation) after detected compared to undetected errors (Nieuwenhuis et al., 2001; Stahl et al., 2020; Wessel et al., 2018; for a review on post-error adjustments see Danielmeier & Ullsperger, 2011).

Notably, response caution was similarly affected by accuracy and confidence across the lifespan. Thus, while metacognitive accuracy was reduced in older age, a neural correlate of confidence magnitude, the P_e/c_ amplitude, and the behavioural adaptations relative to the reported confidence were consistent across the lifespan. This suggests that it is not metacognitive accuracy per se, but rather the perceived confidence that shapes future behaviour: Despite their failure in matching confidence to task performance, older adults seem to be equally able to use internal states of confidence to change future behaviour adaptively.

### 4.5 Limitations and implications

One limitation of the present study is the confined number of trials available for analysis after defining conditions of interest. Due to an unforeseen highly skewed use of the confidence scale, it was impossible to apply a factorial design while retaining four distinct confidence levels. In particular for correct trials, the variance in confidence ratings was low, which is a common problem in metacognition research (for a review, see Wessel, 2012).

A second shortcoming is the number of participants retained for the analyses. When designing the experiment, we tried to find an optimal balance between task difficulty, feasibility for all ages, and gaining many trials while ensuring that especially older adults were not exhausted at the end of the experiment. However, the combination of a substantial number of response alternatives, time pressure, and discriminability of stimuli was demanding, leading to an undesirably large number of participants to be excluded from the analyses.

Nevertheless, our findings provide important insights into ageing effects on metacognition, integrating approaches from error detection and decision confidence research. In contrast to the metacognitive evaluation itself, the effect of confidence on subsequently adapting response caution was well preserved in older adults. Thus, training the metacognitive evaluation of fundamental decisions in older adults might constitute a promising endeavour (and has been shown to work for mathematical problem solving [Pennequin et al., 2010]).

## 5. Conclusion

The study of error detection and confidence in the context of healthy ageing have advanced largely in parallel. Our study demonstrates that confidence shapes our behavioural and neural processing of decisions and should be considered to investigate age-related effects on error processing and metacognitive abilities. Interestingly, the N_e/c_, but not the P_e/c_ amplitude was differentially modulated by confidence across the lifespan, suggesting that the decreasing accuracy of metacognitive judgements with older age might be related to impaired integration of neural correlates of conflict detection and decision confidence.

## Acknowledgements

We thank all colleagues from the Institute of Neuroscience and Medicine (INM-3), Cognitive Neuroscience, and the Decision Neuroscience Lab at the Melbourne School of Psychological Sciences for valuable discussions and their support.

