## Supplemental File for "Neural correlates of metacognition across the adult lifespan"

### Supplement

#### S1. Age distribution and behavioural analysis of the subgroups for EEG analyses

As only participants with a minimum of six trials in each condition were factored into statistical analyses of ERP amplitudes, we formed three subgroups of participants (see main text for more information). We collapsed the confidence ratings to two levels to analyse errors in Subgroup 2, and correct responses in Subgroup 3, respectively. Unfortunately, these two subgroups sacrificed a substantial number of datasets. However, we ensured by inspecting the number of participants per decade that age was still reasonably equally distributed within all subgroups, justifying the examination of the effect of ageing (Figure S1).

To examine how behavioural parameters were affected by the reorganisation of confidence levels, we ran additional analyses for RT and response caution, separating errors and correct responses, and using the same levels of confidence and the same subgroups of participants as for the focussed control analyses of the EEG data. Moreover, we assured that Phi showed the same modulation by age within the subgroups as in the full sample.

***Results***

In the subgroup for errors (Subgroup 1), Phi was still negatively correlated with age ($r(42) = -.474$, $p = .001$). The ANCOVA for RT yielded significant main effects of age ($F(1,42) = 13.213$, $p = .001$, $\eta^2_p = .239$) and confidence ($F(1,42) = 5.077$, $p = .030$, $\eta^2_p = .108$). Errors rated as ‘surely wrong’ were on average faster than errors rated with any higher confidence, but the difference was not significant in a follow-up *t*-test. The ANCOVA for response caution did not yield any significant effects.

In the subgroup for correct trials (Subgroup 3), Phi decreased with age ($r(49) = -.536$, $p < .001$), and RT was also modulated by age ($F(1,49) = 11.583$, $p = .001$, $\eta^2_p = .191$) and confidence ($F(1,49) = 31.378$, $p < .001$, $\eta^2_p = .390$), as revealed in the ANCOVA with the within-subject factor confidence. A follow-up *t*-test showed significantly faster RTs for ‘surely correct’ compared to lower confidence ratings ($t(50) = 12.465$, $p < .001$, $d = 1.744$). Response caution after correct trials did not show any significant effects of age or confidence.

Thus, the reorganisation of trials and confidence levels did not affect the direction of confidence- or age-related effects on behaviour and was consequently used to analyse the EEG data as described.

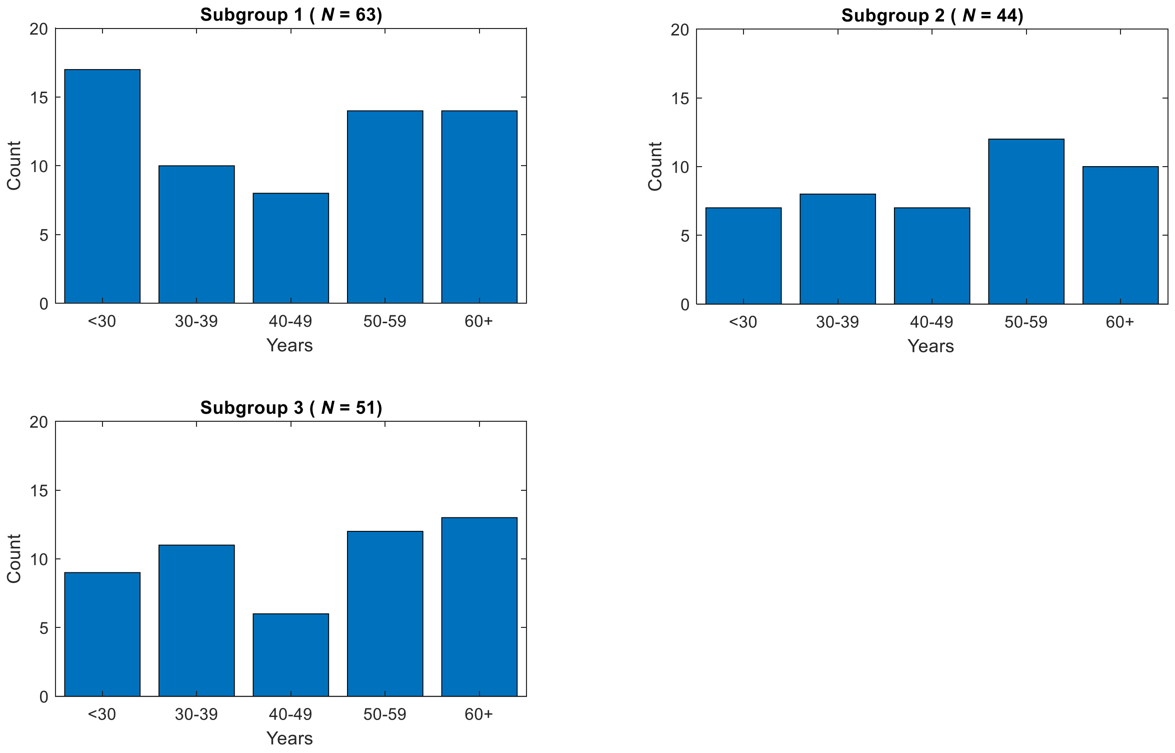


*Figure S1.* Number of participants per decade within the three subgroups used for the analysis of EEG data. Subgroup 1 was used to analyse the effects of accuracy and confidence (3 levels) of all trials on the ERP amplitudes of interest. Subgroup 2 was used to analyse the modulation of the ERPs by confidence (2 levels) for errors and Subgroup 3 for the same analysis for correct responses.

**S2. Modulation of ERPs by confidence, independent of accuracy**

In our main analysis, we performed repeated measures ANCOVAs on all trials with the within-subject factor accuracy and the covariate age for the dependent variables $N_{e/c}$ and $P_{e/c}$, respectively. The amplitudes of both ERPs were larger for errors than correct responses, and decreased with age for errors (trend level for $P_{e/c}$). As both components have further been shown to be sensitive to variations in confidence (Boldt & Yeung, 2015), we additionally computed the $N_{e/c}$ and $P_{e/c}$ amplitudes in relation to reported confidence for errors and correct trials combined (three levels: ‘surely wrong’, ‘unsure’, ‘surely correct’). Here, we provide the results for the ANCOVAs including confidence instead of accuracy as the within-subject factor. Note, that the subgroup for this analysis included 49 subjects and that the proportion of correct and incorrect trials differed across confidence levels.

***Results***

*$N_{e/c}$*. The ANCOVA for the $N_{e/c}$ amplitude with the within-subject factor confidence yielded a main effect of confidence ($F(1,47) = 9.798$, $p < .001$, $\eta^2_p = .173$) as the amplitude increased from trials rated as ‘surely correct’ to ‘unsure’ ($t(48) = -3.038$, $p = .004$, d = 0.434), and from trials rated ‘unsure’ to ‘surely wrong’ ($t(48) = -3.393$, $p = .001$, d = 0.485; ‘surely correct’ vs. ‘surely wrong’: $t(48) = -5.342$, $p < .001$, d = 0.763). Moreover, we found a main effect of age ($F(1,47) = 6.940$, $p = .011$, $\eta^2_p = .129$) and a significant interaction between age and confidence ($F(1,47) = 3.866$, $p = .033$, $\eta^2_p = .076$). Specifically, the $N_{e/c}$ amplitude decreased with age for ‘unsure’ trials ($r(47) = .454$, $p = .001$), but not for ‘surely correct’ trials ($r(47) = .130$, $p = .373$), as confirmed in follow-up correlation analyses. The amplitude of ‘surely wrong’ trials also decreased with age, which did not remain significant after correction for multiple comparisons though ($r(47) = .306$, $p = .033$).

*$P_{e/c}$*. The ANCOVA for the $P_{e/c}$ amplitude with the within-subject factor confidence yielded a main effect of confidence ($F(2,94) = 5.235$, $p = .012$, $\eta^2_p = .100$). Follow-up *t*-tests revealed that

the amplitude on trials rated as 'surely correct' was significantly lower than on trials rated as 'unsure' ($t(48) = 4.507$, $p < .001$, $d = 0.643$) or 'surely wrong' ($t(48) = 5.339$, $p < .001$, $d = 0.763$; 'unsure' vs. 'surely wrong': $t(48) = 1.605$, $p = .115$). However, age did not have an interacting effect on the confidence modulation ($F(2,94) = 1.448$, $p = .241$), and the main effect of age was not significant ($F(1,47) = 1.059$, $p = .309$).

Together, these results replicate previous findings of a confidence-related modulation of the $N_{e/c}$ and $P_{e/c}$ amplitudes (Boldt & Yeung, 2015; Scheffers & Coles, 2000). Moreover, we provide evidence, for the first time, that the $N_{e/c}$, but not the $P_{e/c}$ was differently modulated by ageing across confidence levels. Notably, it has to be considered that the percentage of errors within each confidence level varied substantially between participants and across the lifespan. However, the same holds for the opposite conclusion, that is, a potential modulation of the ERP amplitudes by accuracy is always inherently connected to confidence (e.g., Fleming et al., 2012). Therefore, the more robust analysis, in our opinion, is the separate examination of correct and incorrect trials, which we report in the main article.

Experimental Psychology: Human Perception and Performance, 26(1), 141–151. https://doi.org/10.1037/0096-1523.26.1.141

**S3. Stimulus-related ERPs of conflict processing**

When investigating age-related alterations in neural correlates of response evaluation, the interval between stimulus presentation and response is also informative – in particular the N2 and the P300 components of the ERP. These have been related to error processing as indexing stimulus conflict monitoring (N2) and error-related attention reallocation (P300; Groom & Cragg, 2015; Polich, 2007; Yeung & Cohen, 2006). Research has shown a decline of both components in older age (Korsch et al., 2016; Lucci et al., 2013). Assessing the modulation of these components by age allowed us to draw conclusions about the specificity of potential modulations of the $N_{e/c}$ and $P_{e/c}$ in our metacognitive task. We, therefore, additionally computed the N2 and the P300 components using the stimulus-locked data.

The epochs were cut at 1,500 ms after target stimulus presentation, and the preprocessing was equivalent to the response-locked data. The N2 was quantified as the peak amplitude in the time window from 150 to 300 ms at Cz, and the P300 in the time window from 200 to 500 ms at POz (Groom & Cragg, 2015; Klawohn et al., 2020; Polich, 2007).

***Results***

As expected, a repeated measures ANCOVA for the N2 with the within-subject factor accuracy and the covariate age revealed a main effect of accuracy ($F$(1,61) = 6.729, $p$ = .012, $\eta^2_p$ = .099). Follow-up $t$-tests confirmed larger N2 amplitudes for errors compared to correct responses ($t$(62) = 3.226, $p$ = .002, $d$ = 0.407). Importantly, however, there was no main effect of age or an interaction between accuracy and age. The ANCOVA for the P300 did not show a main effect of accuracy. Similarly to the N2, no effect of age or an interaction between age and accuracy was observed.

This suggests that both the monitoring of stimulus conflict and the attention-related evaluation of conflict were comparable across the lifespan. Therefore, we could infer that age-related differences in early stimulus-related conflict monitoring could not account for subsequent modulations of response processing.

### Table S1

*Distribution of confidence ratings within errors and correct responses across participants.*

| Variable | ANCOVA | | | Follow-up *t*-tests | | | | | | Follow-up correlation analyses | | | |
|---|---|---|---|---|---|---|---|---|---|---|---|---|---|
| | Main effect confidence | Main effect age | Interaction confidence *age | Level 1 vs. level 2 | Level 1 vs. level 3 | Level 1 vs. level 4 | Level 2 vs. level 3 | Level 2 vs. level 4 | Level 3 vs. level 4 | Level 1 | Level 2 | Level 3 | Level 4 |
| Proportion confidence levels (error) | **$F$(3,189) = 28.696, $p$ < .001, $\eta^2_p$ = .313** | - | **$F$(3,189) = 15.722, $p$ < .001, $\eta^2_p$ = .200** | **$t$(64) = 10.780, $p$ < .001, *d = 1.337*** | **$t$(64) = 4.531, $p$ < .001, *d = 0.562*** | **$t$(64) = 2.988, $p$ = .004, $d$ = 0.371** | **$t$(64) = -6.294, $p$ < .001, $d$ = 0.781** | **$t$(64) = -7.844, $p$ < .001, $d$ = 0.973** | $t$(64) = -2.388, $p$ = .020 | **$r$(63) = -.543, $p$ < .001** | $r$(63) = .083, $p$ = .511 | **$r$(63) = .450, $p$ < .001** | $r$(63) = .253, $p$ = .042 |
| Proportion confidence levels (correct) | **$F$(3,189) = 151.589, $p$ < .001, $\eta^2_p$ = .706** | - | **$F$(3,189) = 24.160, $p$ < 001, $\eta^2_p$ = .277** | $t$(64) = -1.505, $p$ = .137 | **$t$(64) = -6.557, $p$ < .001, *d = 0.814*** | **$t$(64) = -32.329, $p$ < .001, $d$ = 4.010** | **$t$(64) = -6.290, $p$ < .001, $d$ = 0.780** | **$t$(64) = -31.286, $p$ < .001, $d$ = 3.880** | **$t$(64) = -15.626, $p$ < .001, $d$ = 1.938** | $r$(63) = .222, $p$ = .076 | $r$(63) = .276, $p$ = .026 | **$r$(63) = .530, $p$ < .001** | **$r$(63) = -.532, $p$ < .001** |

*Notes.* ANCOVAs for the proportion of confidence ratings, separately for errors and correct responses, with the within-subject factor confidence (4 levels; coded from 1: 'surely wrong' to 4: 'surely correct') and the covariate age ($N$ = 65). In case of significant effects, follow-up *t*-tests between confidence levels and/or correlation analyses with age are reported.

### Table S2

*Modulation of behavioural and neural parameters by correctness.*

| Variable | ANCOVA | | | Follow-up *t*-tests | Follow-up correlation analyses | |
|---|---|---|---|---|---|---|
| | Main effect accuracy | Main effect age | Interaction accuracy *age | Error vs. correct | Error | Correct |
| Mean confidence | **$F(1,63) = 164.008$, $p < .001$, $\eta^2_p = .722$** | $F(1,63) = 2.009$, $p = .161$ | **$F(1,63) = 37.433$, $p < .001$, $\eta^2_p = .373$** | **$t(64) = -16.774$, $p < .001$, $d = 2.081$** | **$r(63) = .471$, $p < .001$** | **$r(63) = -.523$, $p < .001$** |
| RT | **$F(1,63) = 4.188$, $p = .045$, $\eta^2_p = .062$** | **$F(1,63) = 18.164$, $p < .001$, $\eta^2_p = .224$** | $F(1,63) = 1.267$, $p = .265$ | **$t(64) = 2.937$, $p = .005$, $d = 0.364$** | - | - |
| Response caution[a] | **$F(1,63) = 6.929$, $p = .011$, $\eta^2_p = .099$** | $F(1,63) = 0.316$, $p = .576$ | $F(1,63) = 2.987$, $p = .089$ | **$t(64) = 2.950$, $p = .004$, $d = 0.357$** | - | - |
| $N_{e/c}$ | **$F(1,61) = 15.209$, $p < .001$, $\eta^2_p = .200$** | **$F(1,61) = 5.999$, $p = .029$, $\eta^2_p = .076$** | **$F(1,61) = 5.999$, $p = .017$, $\eta^2_p = .090$** | **$t(62) = -4.544$, $p < .001$, $d = 0.572$** | **$r(61) = .326$, $p = .009$** | $r(61) = .140$, $p = .272$ |
| $P_{e/c}$ | **$F(1,61) = 23.886$, $p < .001$, $\eta^2_p = .281$** | $F(1,61) = 0.180$, $p = .673$ | **$F(1,61) = 11.836$, $p = .001$, $\eta^2_p = .163$** | **$t(62) = 4.507$, $p < .001$, $d = 0.568$** | $r(61) = -.237$, $p = .061$ | $r(61) = .199$, $p = .118$ |
| N2 | **$F(1,61) = 6.729$, $p = .012$, $\eta^2_p = .099$** | $F(1,61) = 0.633$, $p = .429$ | $F(1,61) = 0.785$, $p = .380$ | **$t(62) = 3.226$, $p = .002$, $d = 0.407$** | - | - |
| P300 | $F(1,61) = 2.383$, $p = .128$ | $F(1,61) = 2.976$, $p = .090$ | $F(1,61) = 0.240$, $p = .626$ | - | - | - |

*Notes.* ANCOVAs for the behavioural and electrophysiological variables including all trials, with the within-subject factor accuracy (2 levels; error, correct) and the covariate age. In case of significant effects, follow-up *t*-tests between error and correct trials and/or correlation analyses with age are reported.

[a] For the measure of response caution, the factor accuracy refers to the previous trial.

**Table S3**

*Modulation of behavioural parameters by confidence.*

| Variable | ANCOVA | | | Follow-up *t*-tests | | | Follow-up correlation analyses | | |
|---|---|---|---|---|---|---|---|---|---|
| | Main effect confidence | Main effect age | Interaction confidence*age | Level 1 vs. level 2 | Level 1 vs. level 3 | Level 2 vs. level 3 | Level 1 | Level 2 | Level 3 |
| ER | **$F(2,124) = 97.426$, $p < .001$, $\eta^2_p = .611$** | $F(1,62) = 2.443$, $p = .128$ | **$F(2,124) = 5.264$, $p = .009$, $\eta^2_p = .078$** | **$t(63) = 19.279$, $p < .001$, $d = 2.410$** | **$t(64) = 35.711$, $p < .001$, $d = 4.429$** | **$t(63) = 10.330$, $p < .001$, $d = 1.291$** | ($r(63) = .037$, $p = .774$ | ($r(63) = -.151$, $p = .229$ | **($r(63) = .568$, $p < .001$** |
| RT | **$F(2,114) = 13.132$, $p < .001$, $\eta^2_p = .187$** | **$F(1,57) = 19.159$, $p < .001$, $\eta^2_p = .252$** | **$F(2,114) = 3.793$, $p = .030$, $\eta^2_p = .062$** | **$t(58) = -5.236$, $p < .001$, $d = 0.682$** | $t(58) = 2.137$, $p = .037$ | **$t(58) = 9.198$, $p < .001$, $d = 1.198$** | **$r(57) = .502$, $p < .001$** | **$r(57) = .567$, $p < .001$** | $r(57) = .261$, $p = .046$ |
| Response caution[a] | **$F(2,110) = 2.897$, $p = .059$, $\eta^2_p = .050$** | $F(1,55) = 0.378$, $p = .541$ | $F(2,110) = 0.901$, $p = .409$ | $t(56) = -0.723$, $p = .473$ | **$t(56) = 3.066$, $p = .003$, $d = 0.406$** | **$t(56) = 3.448$, $p = .001$, $d = 0.457$** | - | - | - |

*Notes.* ANCOVAs for the behavioural variables including all trials, with the within-subject factor confidence (3 levels; coded as 1: 'surely wrong', 2: 'unsure', and 3: 'surely correct') and the covariate age. In case of significant effects, follow-up *t*-tests between confidence levels and/or correlation analyses with age are reported.

[a] For the measure of response caution, the factor confidence refers to the previous trial.

**Table S4**

*Modulation of behavioural and neural parameters by confidence, separately for errors and correct responses.*

| Variable | ANCOVA | | | Follow-up *t*-tests | Follow-up correlation analyses | |
|---|---|---|---|---|---|---|
| | Main effect confidence | Main effect age | Interaction confidence*age | Level 1 vs. level 2 | Level 1 | Level 2 |
| RT (error) | **$F$(1,42) = 5.077, $p$ = .030, $\eta^2_p$ = .108** | **$F$(1,42) = 13.213, $p$ = .001, $\eta^2_p$ = .239** | $F$(1,42) = 3.122, $p$ = .085 | $t$(50) = -1.812, $p$ = .077 | - | - |
| RT (correct) | **$F$(1,49) = 31.378, $p$ < .001, $\eta^2_p$ = .390** | **$F$(1,49) = 11.583, $p$ = .001, $\eta^2_p$ = .191** | $F$(1,49) = 2.738, $p$ = .104 | **$t$(50) = 12.465, $p$ < .001, $d$ = 1.744** | - | - |
| Response caution[a] (error) | $F$(1,42) = 1.283, $p$ = .264 | $F$(1,42) = 0.245, $p$ = .623 | $F$(1,42) = 0.358, $p$ = .553 | - | - | - |
| Response caution[a] (correct) | $F$(1,49) = 1.424, $p$ = .239 | $F$(1,49) = 0.250, $p$ = .619 | $F$(1,49) = 0.741, $p$ = .393 | - | - | - |
| $N_e$ | **$F$(1,42) = 3.461, $p$ = .070, $\eta^2_p$ = .076** | **$F$(1,42) = 10.787, $p$ = .002, $\eta^2_p$ = .204** | $F$(1,42) = 1.447, $p$ = .236 | **$t$(43) = -2.309, $p$ = .026, $d$ = 0.348** | **$r$(42) = .433, $p$ = .003** | **$r$(42) = .350, $p$ = .020** |
| $N_c$ | **$F$(1,49) = 12.624, $p$ = .001, $\eta^2_p$ = .205** | $F$(1,49) = 1.239, $p$ = .271 | **$F$(1,49) = 7.746, $p$ = .008, $\eta^2_p$ = .136** | **$t$(50) = -2.699, $p$ = .009, $d$ = 0.438** | $r$(49) = .278, $p$ = .048 | $r$(49) = -.027, $p$ = .849 |
| $P_e$ | $F$(1,42) = 0.000, $p$ = .996 | $F$(1,42) = 1.235, $p$ = .273 | $F$(1,42) = 0.476, $p$ = .494 | *[exploratory]* **$t$(43) = 2.157, $p$ = .037, $d$ = 0.325** | - | - |
| $P_c$ | **$F$(1,49) = 5.065, $p$ = .029, $\eta^2_p$ = .094** | $F$(1,49) = 0.006, $p$ = .939 | $F$(1,49) = 0.186, $p$ = .668 | **$t$(50) = 5.838, $p$ < .001, $d$ = 0.818** | - | - |

*Notes.* ANCOVAs for the behavioural and electrophysiological variables for the subgroups of the focussed control analyses, separately for errors and correct responses. The within-subject factor confidence is collapsed to two levels (errors: coded as 1: 'surely wrong' and 2: 'higher confidence'; correct trials: coded as 1: 'lower confidence', 2: 'surely correct') and the covariate of interest is age. In case of significant effects, follow-up *t*-tests between confidence levels and/or correlation analyses with age are reported.

[a] For the measure of response caution, the factor confidence and the distinction error/correct refer to the previous trial.
